# Cell-Type Selective Markers Represented in Whole-Kidney RNA-Seq Data

**DOI:** 10.1101/348615

**Authors:** Jevin Z. Clark, Lihe Chen, Chung-Lin Chou, Hyun Jun Jung, Jae Wook Lee, Mark A. Knepper

## Abstract

Bulk-tissue RNA-Seq is seeing increasing use in the study of physiological and pathophysiological processes in the kidney. However, the presence of multiple cell types in kidney complicates the data interpretation. Here we address the question, “What cell types are represented in whole-kidney RNA-Seq data?” to identify circumstances in which bulk-kidney RNA-Seq can be successfully interpreted. We carried out RNA-Seq in mouse whole kidneys and microdissected renal tubule segments. To aid in the interpretation of the data, we compiled a database of cell-type selective protein markers for 43 cell types believed to be present in kidney tissue. The whole-kidney RNA-Seq analysis identified transcripts corresponding to 17742 genes, distributed over 5 orders of magnitude of expression level. Markers for all 43 curated cell types were detectable. Analysis of the cellular makeup of mouse and rat kidney, calculated from published literature, suggests that proximal tubule cells account for more than half of the mRNA in a kidney. Comparison of RNA-Seq data from microdissected proximal tubules with whole-kidney data supports this view. RNA-Seq data for cell-type selective markers in bulk-kidney samples provide a valid means to identify changes in minority-cell abundances in kidney tissue. Because proximal tubules make up a substantial fraction of whole-kidney samples, changes in proximal tubule gene expression can be assessed presumptively by bulk-kidney RNA-Seq, although results could potentially be obscured by the presence of mRNA from other cell types. The dominance of proximal tubule cells in whole-kidney samples also has implications for the interpretation of single-cell RNA-Seq data.

## INTRODUCTION

RNA-Seq is a method for identifying and quantifying all mRNA species (considered in this paper) in a sample as well as many non-coding RNA species.^1, 2, 3^ Like RT-PCR, the first step of RNA-Seq is reverse transcription of all mRNAs to give corresponding cDNAs. However, unlike RT-PCR, which amplifies only one cDNA target, RNA-Seq amplifies all cDNAs in the sample through use of adaptors that are ligated to the ends of each cDNA.^4^ The read-out for RNA-Seq employs next-generation DNA sequencers to identify specific sequences that map to each mRNA transcript coded by the genome of a particular species (the ‘transcriptome’). This allows counting of the number of ‘reads’ for each transcript as a measure of the total amount of each transcript in the original sample. So, RNA-Seq can be viewed simplistically like quantitative RT-PCR, but more expansive and unbiased.^1^ The abundance of a given transcript is assumed to be proportional to the number of independent sequence ‘reads’ normalized to the annotated exon length of each individual gene and to the total reads obtained for a sample. This calculation yields transcripts per million or ‘TPM’ as termed in this paper.^5^

RNA-Seq has seen increased use in recent years, in part because of the ease of execution and the availability of next-generation DNA sequencers.^6^ Because of the existence of private-sector biotechnology companies, even small laboratories can successfully carry out RNA-Seq studies in lieu of quantitative RT-PCR. Many recent reports using RNA-Seq employ “bulk-tissue RNA-Seq” in which complex tissues containing multiple cell types are analyzed. The limitation of this approach is that it is usually impossible to determine which cell types in the mixture are responsible for observed changes in mRNA abundances. Furthermore, strong responses in minority cell types may be masked by a lack of response in more abundant cell types.^7^ Similar limitations apply to other analytical modalities, such as proteomics.

A solution to this problem in kidney is to isolate specific cell types using renal tubule micro-dissection prior to small sample RNA-Seq as described by Lee et al.^8, 9^ All 14 renal tubule segments plus glomeruli have been profiled in this way. In structures that contain more than one cell type, transcriptomes of each cell type can be determined using single-cell RNA seq (scRNA-Seq).^10, 11, 12, 13, 14, 15, 16, 17^ However, RNA-Seq in single tubules or single cells is not always feasible, e.g. in pathophysiological models or biopsy samples when inflammation or fibrosis limits tissue dissection or single-cell dissociation. In this context, we ask the question, “Despite the existence of multiple cell types in bulk-kidney samples, what information about specific cell types can be gleaned from whole-kidney RNA-Seq?”

## RESULTS

### What mRNA species are detectable in whole-kidney RNA-Seq analysis?

We carried out RNA-Seq analysis in three whole-kidney samples from untreated 2-month-old male C57BL/6 mice. Supplemental Figure 1 shows that the percentage of uniquely mapped reads exceeded 85% of the total reads indicating high data quality for all three samples. Total reads for each of the three samples exceeded 66 million reads. Figure 1 shows the reads that mapped to selected genes expressed over a broad range of TPM levels. It can be seen that faithful, selective mapping to exons was obtained down to a TPM value of about 0.15 in this study, or an expression rank of 17742. For example, the reads for Oxtr, coding for the oxytocin receptor (TPM=0.15), thought to be expressed selectively in macula densa cells,^18^ are clearly mapped only to exons of the Oxtr gene indicating the specificity of the measurement for spliced Oxtr mRNA (see Supplemental Dataset 1 for mapping of reads for other transcripts with TPM around 0.15). In contrast, exon-specific mapping is ambiguous for Epo, the transcript that codes for erythropoietin (TPM=0.09). Overall, we conclude that 17742 transcripts out of approximately 21000 protein-coding genes in the mouse genome can be detected and quantified in whole kidney samples with the technical approach used here. The whole-kidney TPM values for all transcripts down to rank 17742 are presented at a publicly accessible webpage (https://hpcwebapps.cit.nih.gov/ESBL/Database/MouseWK/) and as Supplemental Dataset 2. Mapping of whole kidney RNA-Seq reads on a genome browser can be viewed by clicking on “UCSC Genome Browser” at this site. Since the data in this paper were obtained exclusively from 2-month-old male C57BL/6 mice, the reader is cautioned about possible differences that may occur on the basis of gender, age, mouse strain, animal species, food intake, etc. Further studies will be needed to identify the effects of these variables.

**Figure 1.**
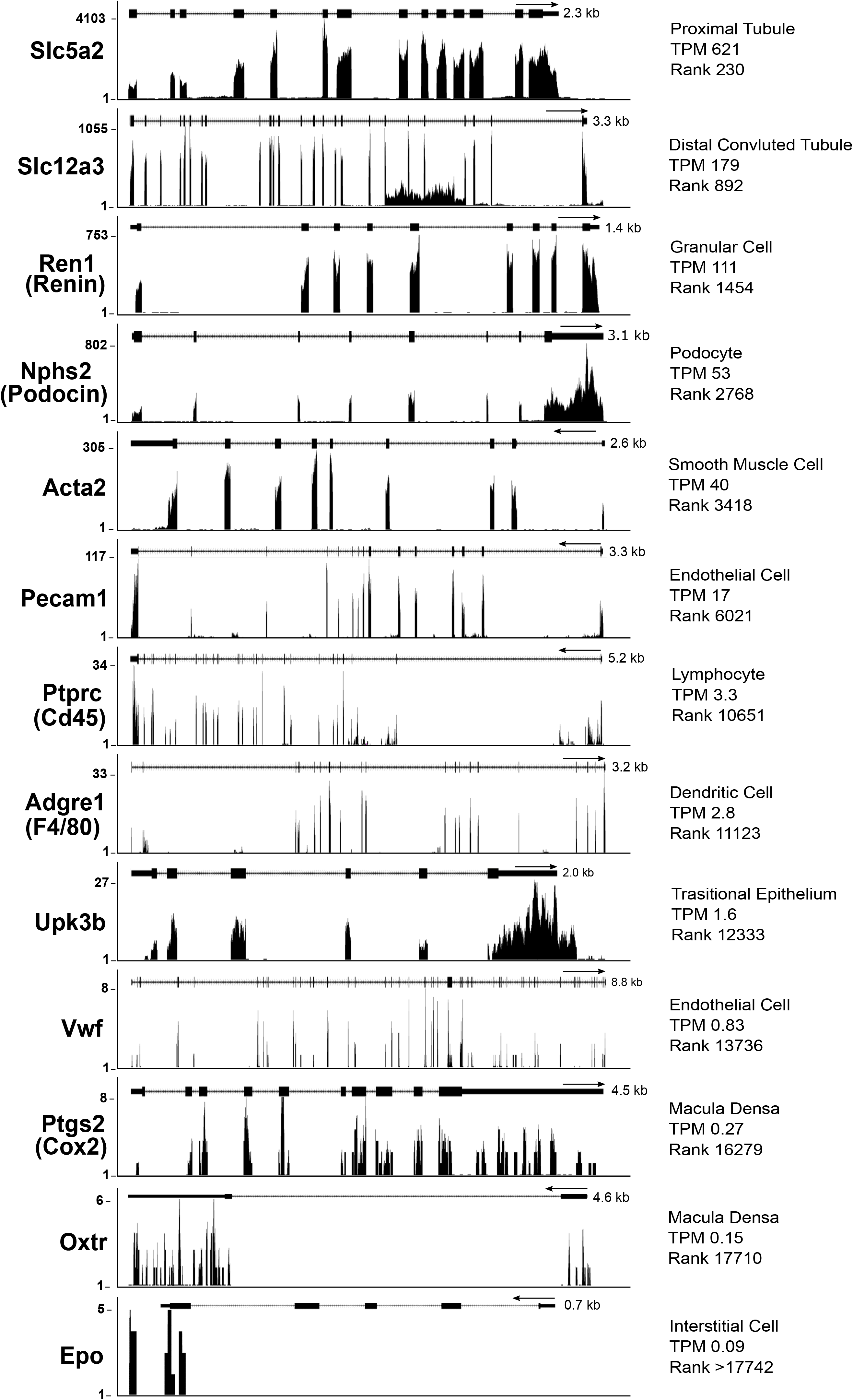
Visualization of the RNA-Seq reads for representative transcripts. Cell-type selective genes from indicated cell types with their mRNA length, TPM, and Rank values. Genes with TPM greater than 0.15 are within a confident detectable range. Data were visualized in the UCSC Genome Browser. Vertical axis shows read counts. Map of exon/intron organization of each gene is shown on top of individual panels.

### What cell types are represented in whole-kidney RNA-Seq data?

Based on a variety of data types (Methods), we curated a list of 43 cell types that are thought to exist in the kidney and representative protein markers that have been claimed to be specific to or selective for these cell types. The cell types, the markers and whole kidney TPM values for mRNAs corresponding to the markers are presented in Supplemental Dataset 3 and at a permanent, publicly available webpage (https://hpcwebapps.cit.nih.gov/ESBL/Database/MouseWK/WKMarkers.html). Selected values are presented in Tables 1 and 2. Table 1 shows TPM values for selected markers of epithelial cell types and Table 2 shows TPM values for selected markers of non-epithelial cell types. As seen in Table 1, markers for each epithelial cell type are highly expressed with the exception of macula densa cells. The TPM values for many non-epithelial cell type markers are above the TPM=0.15 threshold defined above (Table 2 and Supplemental Dataset 3). Overall, based on the markers that we have curated, we conclude that mRNAs from at least 43 cell types are detectable in whole kidney RNA-Seq samples from mouse. This includes various blood-borne cells, stromal cells and endothelial cells.

**Table 1.** Selected markers for renal epithelial cells in mouse whole kidney with corresponding TPM and Rank values. The full marker dataset values are listed in Supplemental Dataset 3.

| Cell Type | Gene Symbol | Common Name | TPM | Rank |
| --- | --- | --- | --- | --- |
| Podocyte | **Nphs2** | Podocin | 53.3 | 2768 |
| Proximal (S1) | **Slc5a2** | Type 2 Na-glucose cotransporter (SGLT2) | 621.2 | 230 |
| Thin Ascending Limb | **Clcnka** | Chloride channel, voltage sensitive, kidney type A | 60.4 | 2511 |
| Thick Ascending Limb | **Slc12a1** | Type 2 Na-K-2Cl cotransporter (NKCC2) | 333.8 | 470 |
| Macula Densa | **Ptgs2** | Prostaglandin-endoperoxide synthase 2 (COX2) | 0.3 | 16278 |
| Distal Convoluted Tubule | **Slc12a3** | Thiazide-sensitive Na-Cl cotransporter (NCC) | 179.1 | 892 |
| Connecting Tubule | **Calb1** | Calbindin 1 | 316.1 | 499 |
| Principal Cell | **Aqp2** | Aquaporin-2 | 464.1 | 317 |
| Intercalated Cell, Type A | **Slc4a1** | Chloride-bicarbonate transporter 1 (AE1) | 17.4 | 5905 |
| Intercalated Cell, Type B | **Slc26a4** | Pendrin | 39.6 | 3484 |
| Inner Medullary Collecting Duct Cell | **Slc14a2** | Urea channel, epithelial | 20.1 | 5484 |
| Transitional Epithelium | **Upk1a** | Uroplakin 1a | 7.4 | 8472 |

**Table 2.** Selected markers for renal non-epithelial cells in mouse whole kidney with corresponding TPM and Rank values. The full marker dataset values are listed in Supplemental Dataset 3.

| Cell Type | Gene Symbol | Common Name | TPM | Rank |
| --- | --- | --- | --- | --- |
| Basophil | **Cd69** | Cd69 antigen | 0.2 | 16835 |
| B-Lymphocyte (follicular) | **Cd22** | B-cell receptor | 0.2 | 17615 |
| Dendritic Cell | **Adgre1** | Adhesion G protein-coupled receptor E1 (F4/80) | 2.7 | 11123 |
| Endothelial Cell | **Pecam1** | Platelet/endothelial cell adhesion molecule 1 | 16.8 | 6021 |
| Fibroblast | **Pdgfrb** | Platelet derived growth factor receptor, beta | 7.8 | 8347 |
| Granular Cell of Afferent Arteriole | **Ren1** | Renin 1 | 111.3 | 1454 |
| Macrophage | **Cd68** | Macrosialin | 4.9 | 9629 |
| Monocyte | **Cd14** | Monocyte differentiation antigen CD14 | 5.3 | 9436 |
| Neuronal Cell (Axon Only) | **Stx1a** | Syntaxin 1A (brain) | 0.5 | 14797 |
| Smooth Muscle Cell | **Acta2** | Actin, alpha 2, smooth muscle | 40.5 | 3418 |
| Polymorphonuclear Leukocyte | **Csf3r** | colony stimulating factor 3 receptor (granulocyte) | 0.2 | 16597 |
| T-lymphocyte | **Cd4** | T-cell surface glycoprotein CD4 | 0.5 | 14893 |

### How much do various kidney tubule cell types contribute to TPM values?

Table 3 shows an accounting of the relative contributions of various renal epithelial cell types to the total makeup of the rat and mouse renal tubule in terms of cell number and protein mass. The estimates for rat and mouse were established by integrating several data sources relevant to quantitative renal anatomy.^19, 20, 21, 22^ Full calculations and data sources are available in Supplemental Dataset 4. Values for percentages of cells and protein mass for individual cell types are very similar for mouse and rat and we concentrate on rat values here. Proximal tubule cells account for roughly 52% of the estimated 206 million tubule epithelial cells per kidney. However, they account for approximately 69% of total tubule protein mass, by virtue of their large size compared to other renal tubule cells (Table 3). The second largest contribution is from the thick ascending limb of Henle, contributing 17% of cells and 12% of total protein (Table 3). If mRNA levels parallel protein levels, the contribution of proximal tubules to total mRNA in the renal tubule is also likely to be considerably greater than 50%.

**Table 3.** Contributions of epithelial cell types to whole kidney cell count and mass in rat and mouse.

| Segment/Cell type<sup>a</sup> | Total Cells per Kidney (millions) |  | Percent of Total Cells |  | Total Protein Mass (μg) |  | Percent of Total Protein Mass |  |
| --- | --- | --- | --- | --- | --- | --- | --- | --- |
|  | Rat | Mouse | Rat | Mouse | Rat | Mouse | Rat | Mouse |
| S1 Proximal | 48.36 | 10.19 | 23.52 | 20.08 | 33031 | 8189 | 31.23 | 29.80 |
| S2 Proximal | 48.36 | 10.19 | 23.52 | 20.08 | 33031 | 8189 | 31.23 | 29.80 |
| S3 Proximal | 10.75 | 2.26 | 5.23 | 4.46 | 7340 | 1820 | 6.94 | 6.62 |
| tDL - type 1 | 4.05 | 1.17 | 1.97 | 2.30 | 1497 | 432 | 1.42 | 1.57 |
| tDL - type 2 | 3.31 | 0.44 | 1.61 | 0.86 | 815 | 108 | 0.77 | 0.39 |
| tDL - type 3 | 1.81 | 0.73 | 0.88 | 1.43 | 537 | 215 | 0.51 | 0.78 |
| tAL | 3.00 | 0.87 | 1.46 | 1.72 | 741 | 215 | 0.70 | 0.78 |
| MTAL | 17.73 | 7.55 | 8.62 | 14.87 | 7125 | 3033 | 6.74 | 11.04 |
| cTAL | 17.77 | 3.61 | 8.64 | 7.12 | 5103 | 1038 | 4.82 | 3.78 |
| DCT | 19.90 | 3.78 | 9.68 | 7.45 | 8459 | 1607 | 8.00 | 5.85 |
| CNT Cell | 6.99 | 2.66 | 3.40 | 5.24 | 2011 | 764 | 1.90 | 2.78 |
| CNT A-IC | 1.17 | 0.44 | 0.57 | 0.87 | 335 | 127 | 0.32 | 0.46 |
| CNT B-IC | 3.50 | 1.33 | 1.70 | 2.62 | 1005 | 382 | 0.95 | 1.39 |
| CCD PC | 4.15 | 1.31 | 2.02 | 2.57 | 881 | 277 | 0.83 | 1.01 |
| CCD B-IC | 1.58 | 0.50 | 0.77 | 0.98 | 334 | 105 | 0.32 | 0.38 |
| CCD A-IC | 1.43 | 0.45 | 0.70 | 0.89 | 304 | 96 | 0.29 | 0.35 |
| OMCD - PC | 3.99 | 1.21 | 1.94 | 2.39 | 1013 | 308 | 0.96 | 1.12 |
| OMCD - A-IC | 2.50 | 0.76 | 1.22 | 1.50 | 635 | 193 | 0.60 | 0.70 |
| OMCD - B-IC | 0.16 | 0.05 | 0.08 | 0.10 | 41 | 12 | 0.04 | 0.04 |
| IMCD | 5.15 | 1.25 | 2.50 | 2.46 | 1544 | 374 | 1.46 | 1.36 |
| SUM | 205.63 | 50.74 | -- | -- | 105782.02 | 27484.00 | -- | -- |

a abbreviation, definition; A-IC, Type A Intercalated Cells; B-IC, Type B Intercalated Cells; PC, Primary Cells; PT, Proximal Tubule; tDL, thin descending limb; tAL, thin ascending limb of the loop of Henle; mTAL, medullary thick ascending limb of the loop of Henle; cTAL, cortical thick ascending limb of the loop of Henle; DCT, distal convoluted tubule; CNT, connecting tubule; CCD, cortical collecting duct; OMCD, outer medullary collecting duct; IMC, inner medullary collecting duct
The following sources were used for the calculations: Sperber<sup>22</sup>, Murawski<sup>19</sup>, Knepper<sup>34</sup>, Garg<sup>20</sup>, Guder<sup>21</sup>. All of the calculations and sources are also available in Supplemental Dataset 4.

Wiggins et al. have quantified the cell types that make up the glomerulus in rats,^23^ yielding a median value of 133 podocytes per glomerulus. In each rat kidney, there are 38000 glomeruli per rat kidney X 133 podocytes per glomerulus = 5.1 X 10^6^ podocytes per rat kidney. This value is about 2.4% of the total number of epithelial cells in rat (Table 3). In Bertram et al. a somewhat larger estimate of the number of podocytes per rat glomerulus was obtained (about 181 per glomerulus) which would predict that podocytes make up 3.4% of total epithelial cells (Table 3).^24^ The number of podocytes per mouse kidney is smaller (about 75 per glomerulus).^25^ This would give 20220 glomeruli per mouse kidney X 75 podocytes per glomerulus = 1.5 X 10^6^ podocytes per mouse kidney. This comes out to 3.0% of total epithelial cells in mouse kidney (Table 3). Thus, changes in podocyte transcripts are unlikely to be readily detectable or quantifiable in whole-kidney samples, unless they are specific to the glomerulus. Qiu et al. have described an effective means of obviating this limitation, viz. separate analysis of glomeruli microdissected from kidney samples.^26^

### What fraction of mouse whole kidney mRNA is derived from proximal tubule cells, thick ascending limb cells and collecting duct principal cells?

Because the proximal tubule makes such a large contribution to total epithelial cell number and protein mass (Table 3), it seems possible that whole kidney RNA-Seq measurements could be used as a surrogate for measurements of transcript levels in the proximal tubule. In order to compare the mouse whole kidney transcriptome with that of the mouse proximal tubule, we carried out RNA-Seq in microdissected S2 proximal tubules, manually dissected from the opposite (left) kidney from the one used for whole kidney RNA-Seq analysis. The S2 segment was chosen, rather than S1 or S3, because it is rapidly dissectible without collagenase treatment and clearly identifiable because of its presence in the cortical medullary rays. The S2 proximal data mapped to a total of 18767 genes with mean TPM values greater than 0.1 among the three animals. All of the 12 S2 proximal samples (4 replicates per kidney) had a percent of mapped reads greater than 85, consistent with high data quality (Supplemental Figure 2). The mean TPM values are provided as a publicly accessible web page at https://hpcwebapps.cit.nih.gov/ESBL/Database/MusRNA-Seq/index.html. Figure 2A and 2B show plots of the base 2 logarithms of the whole kidney (WK) versus proximal S2 TPM values for housekeeping and nonhousekeeping genes, respectively. The list of housekeeping genes was taken from Lee et al.^8^ The ratios for all genes were normalized such that the average WK/S2 TPM ratio is 1 for housekeeping genes that have TPM greater than 1. A tight correlation was seen for housekeeping transcripts (Figure 2A). As expected, WK/S2 ratios varied over a broad range for nonhousekeeping transcripts. The lower bound is seen at a ratio of about 0.25 and coincides with the location of S2-specific transcripts, e.g. Slc22a7 and Slc22a13, which mediate organic anion and organic cation secretion, respectively, key functions of the S2 segment.^27^ This suggests that the S2 segment accounts for approximately one quarter of whole kidney mRNA. Kap, a proximal tubule marker expressed in all three subsegments (S1, S2, and S3) is found near the 0.5 ratio line, suggesting that the proximal tubule may account for roughly 50% of whole kidney mRNA.

**Figure 2.**
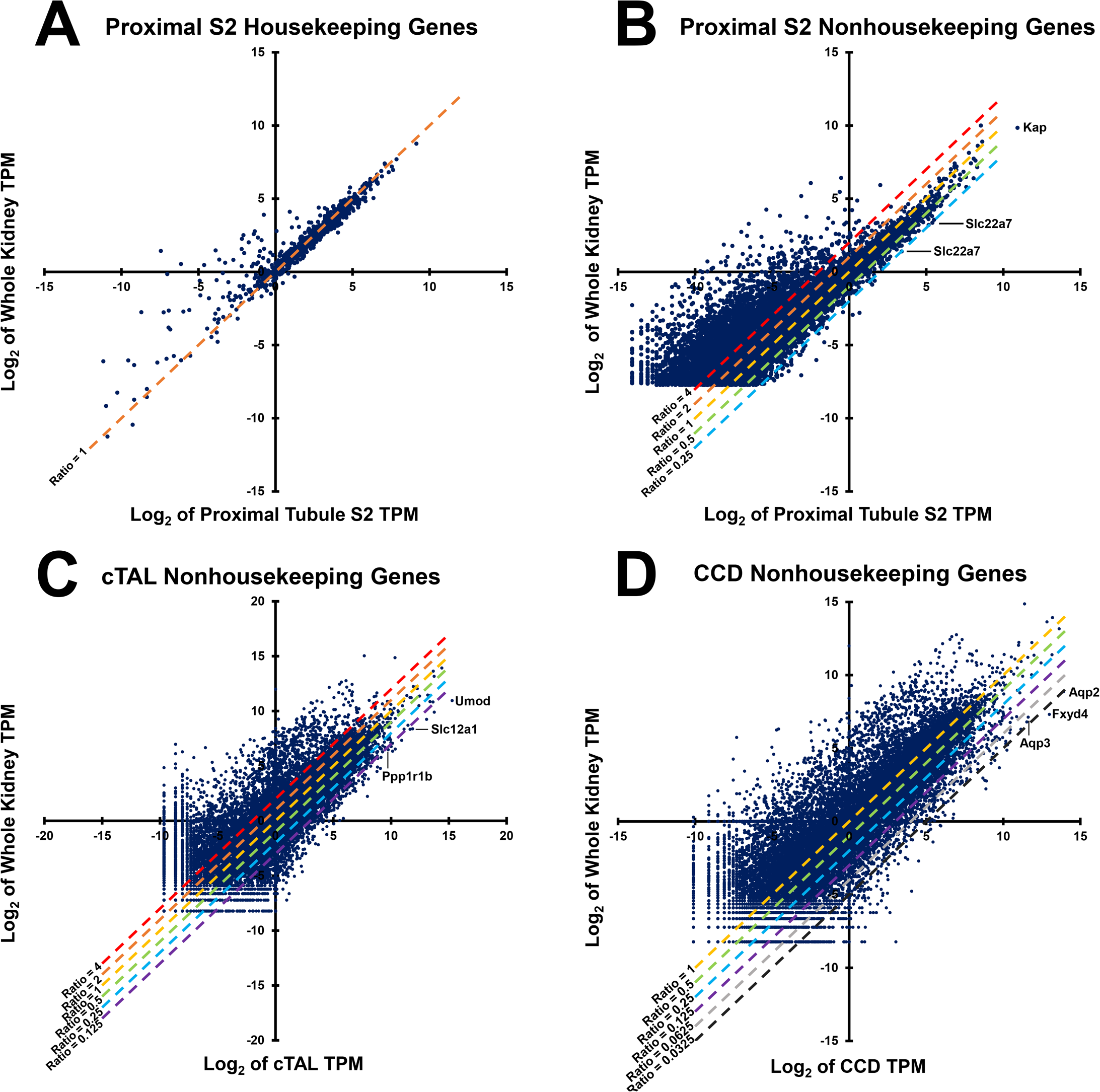
Correlation between whole kidney RNA-Seq and microdissected single-tubule RNA-Seq. (A) Housekeeping genes were plotted for whole-kidney RNA-Seq versus microdissected proximal tubule S2 RNA-Seq. (B-D) Nonhousekeeping genes were plotted for whole kidney RNA-Seq versus the indicated microdissected single tubule RNA-Seq. The dashed lines represent the whole-kidney versus respective tubule RNA-Seq ratios. For (B), each dot is an individual transcript with TPM greater than 0.15. Data are log_2_-transformed before plotting.

TPM values for microdissected mouse cortical thick ascending limbs (cTALs) and cortical collecting ducts (CCDs) were mined from a prior study^10^ and compared to the whole kidney RNA-Seq data from this paper (Figures 2C-D). The lower bound of values for cTAL corresponds to known thick ascending limb markers (Umod, Slc12a1 and Ppp1r1b) just below the ratio 1:8 line. The specific ratios for these markers give an estimate that thick ascending limbs account for roughly 8.8 percent of the total kidney mRNA. This contrasts with a value of about 15 percent based on morphometric analysis in mouse (mTAL plus cTAL) (Table 3), possibly due to dilution of the whole-kidney values by non-epithelial cells not accounted for in the morphometric analysis. The lower bound for CCD cells corresponds to known principal cell markers (Aqp2, Aqp3 and Fxyd4) at a ratio of around 1-to-32, suggesting that principal cells account for around 3 percent of the whole kidney transcriptome.

### What is the contribution of non-epithelial cell types to the overall bulk kidney transcriptome?

Given the estimates of the percent contribution of each epithelial cell type in Table 3 and RNA-Seq data from microdissected tubules from rat kidney,^8^ it is possible to calculate a ‘reconstructed’ bulk kidney transcriptome. This can be compared to rat whole-kidney RNA-Seq data from our laboratory (Gene Expression Omnibus, number GSE70012). The difference between the two can be attributed to non-renal tubule cell types and is presented in Table 4 and Supplemental Dataset 5 in the form of measured:reconstructed ratios. As seen in Table 4, this analysis in rat confirms the presence of several non-renal tubule cell types in bulk kidney tissue and establishes the listed markers as detectible in normal rat kidneys.

**Table 4.**
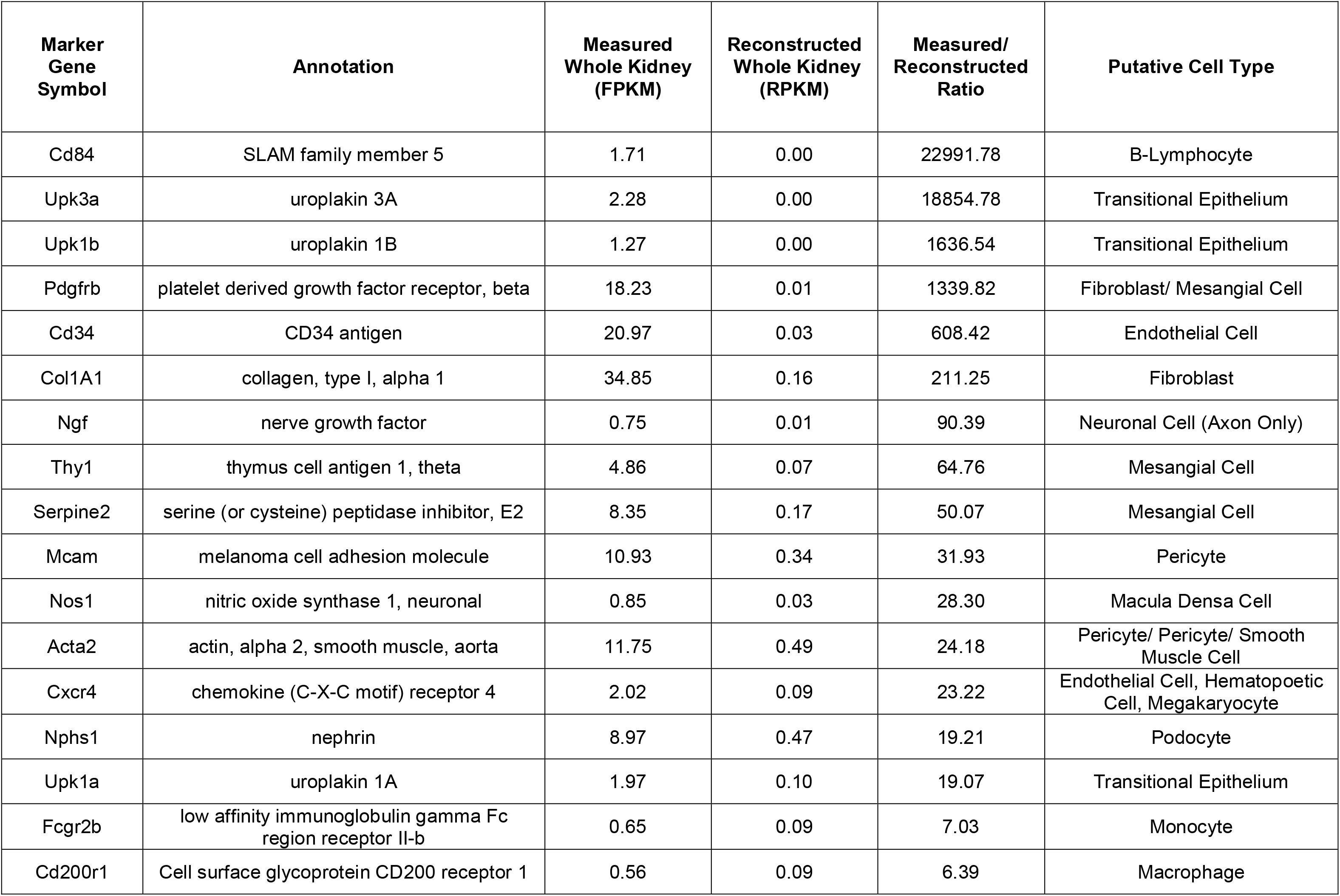
Transcripts highly expressed in rat whole kidney but not in renal tubule epithelia. “Reconstructed Whole Kidney” refers to whole kidney gene expression calculated from rat single tubule RNA-Seq and estimates of percent contribution of each renal tubule cell type. “Measured Whole Kidney” refers to whole kidney RNA-Seq in rats. Cell types correspond to those annotated in Supplemental Dataset 3.

### Reconstructed RNA-Seq transcriptome of whole kidney from scRNA-Seq data

Recently, there have been several reports that provide single-cell RNA-Seq data for many of the known renal tubule cell types.^10, 11, 12, 13, 14, 15, 16, 17^ In theory, single-cell transcriptomes could be used to produce reconstructed bulk-kidney transcriptomes in a manner similar to that presented in the previous section using data from microdissected renal tubules. However, the calculation requires comprehensive transcriptomes in each cell, i.e. a full accounting of the abundances of all expressed transcripts, which appears to correspond to 7000-8000 expressed genes in each cell type.^8^ Figure 3A shows the average number of transcripts quantified in selected individual cell types in a recent scRNA-Seq profiling study that used a state-of-the-art droplet-based method.^15^ Similar values (not shown) were obtained from another recent droplet-based scRNA-Seq studies of kidney.^11^ As can be seen, the average number of transcripts quantified was in the range 274-476. Thus, although the most abundant transcripts were found, the transcriptome list does not appear to be comprehensive despite the use of state-of-the-art methodology. Furthermore, information about gene expression that can identify a particular cell type is conveyed only in nonhousekeeping genes, which constituted less than a third of the total. As shown in Figure 3B, RPKM or TPM values from comprehensive transcriptomic data sets shows that the percent nonhousekeeping transcripts increases beyond that obtained in droplet-based scRNA-Seq of kidney (shaded region). Thus, a goal for the future is to increase the depth of scRNA-Seq transcriptomic analysis for all major cell types in the kidney. A strategy for doing this is proposed in the *Discussion*.

**Figure 3.**
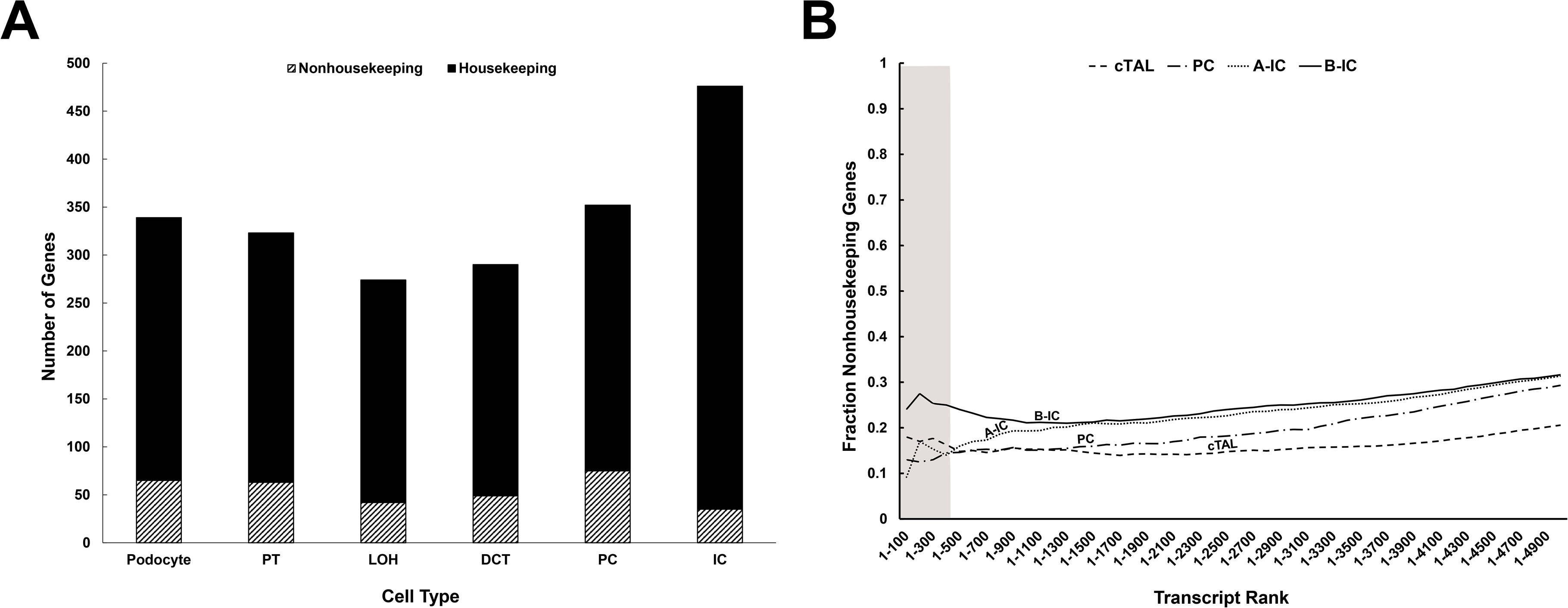
Sequencing depth in single-cell RNA-Seq. (A) Average number of transcripts quantified in selected individual cell types from Park et al. The genes selected had mean transcript count greater than 1 and were categorized into housekeeping and nonhousekeeping genes. The list of housekeeping genes was taken from Lee et al. (B) The cumulative percentage of nonhousekeeping genes are plotted versus TPM rank for mouse whole kidney transcriptome data presented in this paper. The shaded region correlates to the maximum number of transcripts (476) in single cell data as identified in (A).

## DISCUSSION

In this paper, we asked the question, “What information about specific cell types can be gleaned from whole-kidney RNA-Seq?”. To address this, we carried out RNA-Seq analysis of whole mouse kidney samples, yielding a database of 17742 transcripts with TPM values above a threshold of 0.15, determined from examination of mapped reads for a variety of transcripts spanning TPM values from 0.10 to 621 (see Figure 1 and Supplemental Dataset 2). A full report of TPM values for all 17742 transcripts is given at a publicly accessible website. To identify cell types represented in these data, we compiled a list from literature of selective markers for 43 cell types likely present in kidney tissue. These are listed in Supplemental Dataset 3. (Note that we made no attempt to make the marker list totally comprehensive. Readers are encouraged to look up other transcripts of interest at the website of RNA-Seq data: https://hpcwebapps.cit.nih.gov/ESBL/Database/MouseWK/index.html). We detected markers for all 43 cell types, many of them presumably rare in the overall cell count for the kidney. Thus, even for rare cell types, bulk RNA-Seq data can be used to draw inferences about the abundance of a particular cell type or regulation of its marker. For example, an inflammatory process in the kidney is likely to be associated with increases in markers for macrophages (e.g. Adgre1 [F4/80] or Cd68) in whole-kidney RNA-Seq data. Similarly, an increase in mRNA for renin in the kidney may be seen if either the number of afferent arteriolar granular cells increases or when the transcription of the renin gene is increased, both of which have been observed.^28^

Our analysis of the abundances of individual epithelial cell types confirms that proximal tubule cells account for a large fraction of the total kidney substance, most likely at least 50%. The S2 segment alone appears to account for approximately 25% of whole kidney mRNA (Figure 2B). This raises the question of whether whole kidney measurements suffice to assess changes in the proximal tubule. Clearly, changes in proximal tubule mRNA abundance for a particular gene should be detectable in whole kidney samples, although the magnitude of changes will be attenuated by dilution by other cell types. The main problem with interpreting whole kidney changes as tantamount to changes in the proximal tubule is that large changes that are specific to other segments would also be manifest in whole kidney samples. Furthermore, changes in the proximal tubule could be masked by opposite changes in other cell types. Consequently, we do not recommend using whole-kidney or bulk-tissue RNA-Seq as the sole methodology to address hypotheses about the proximal tubule. One approach that may be better in this setting is single-tubule RNA-Seq,^8^ in which proximal tubules are first microdissected from the kidney and then subjected to small sample RNA-Seq analysis. In this paper, we present new single-tubule RNA-Seq data on the transcriptome of microdissected S2 proximal straight tubules and present a comparison with the whole-kidney RNA-Seq data.

The compendium of cell-type selective protein markers provided in this paper is a resource that may be useful to investigators. We caution that the list is not necessarily comprehensive. The list includes multiple markers that have been claimed for certain cell types, many of which were chosen because the protein is present on the cell surface allowing cell sorting. The imprecise definition of the term “cell marker” may lead to uncertainty when interpreting different types of data, thus cell surface markers could be suboptimal for interpretation of RNA-Seq data. Furthermore, many markers have been claimed to be cell-type specific in several cell types, contradicting the specificity claim. In general, we believe that there is a need for a kidney-community oriented effort to define the best cell markers for various uses.

In this paper, we have shown that it is possible to create a ‘reconstructed’ whole-kidney transcriptome from transcriptomes of individual renal tubule segments and information about the relative abundances of each cell type in the kidney from morphometric data. Success with this exercise has helped to validate the accuracy of quantitative RNA-Seq data from structures isolated from the kidney. This bodes well also for establishing the validity of scRNA-Seq measurements.^10, 11, 12, 13, 14, 15, 16, 17^ However, we could not carry out whole-kidney reconstructions using the state-of-the-art scRNA-Seq data that is currently available because the number of transcripts measured in these studies (274-476) fell short of the full depth of cellular transcriptomes (at least 7000-8000).^8^ Thus, although the scRNA-Seq data that have been published represents a very large step forward, there remains an un-reached objective, viz. to push the method so that the scRNA-Seq identifies full transcriptomes for all of the major cell types. Until now, comprehensive scRNA-Seq studies have employed a shotgun approach which involved digestion of the whole kidney and sequencing to obtain transcriptomes for all single cells obtained.^11, 15^ A limitation of this approach is that, as confirmed in this study, proximal tubule cells are much more abundant than any other cell type in the kidney. Consequently, an unbiased sequencing of all cells results in most of the sequencing resources being devoted to proximal tubule cell transcriptomes. As a result, if investigators increase the amount of sequencing to obtain deeper transcriptomes with a shotgun approach, most of the additional effort will be wasted on proximal-tubule cells. To avoid this inefficiency, in the quest to obtain deep transcriptomes in minority cell types, it may be necessary to use microdissection, biochemical procedures, or flow sorting to isolate or enrich those cell types. Already, scRNA-Seq studies have been reported using this strategy for components of the glomerulus^12^ and the collecting duct.^10^

Beyond this reconstruction approach, there is potential value in being able to work in the opposite direction to ‘deconvolute’ bulk-tissue data,^29^ e.g. in the analysis of formalin-fixed paraffin-embedded kidney biopsy samples,^30, 31^ to ascertain what cell types are present in the samples and how they are altered by disease processes. This can succeed qualitatively by identifying cell-type specific transcripts that differ in abundance in a patient sample versus some appropriate reference. However, a difference in a particular transcript could be due either to a change in the number of cells or a change in the expression of the marker in each cell. The use of multiple markers may help to resolve this ambiguity. In the long term, machine learning techniques can be used to generate classifiers from bulk RNA-Seq data that can identify disease processes.^32^

### Summary

RNA-Sequencing (RNA-Seq) is seeing increasing use to assess gene expression in the kidney. To discover pathophysiological mechanisms in animal models of kidney disease, RNA-Seq is often carried out in bulk kidney tissue, consisting of multiple cell types. This study analyzes RNA-Seq data from whole kidneys from normal mice and rats to identify the cell types represented in the data. Markers for 43 different cell types were clearly detectible including all epithelial cell types plus multiple types of vascular cells, stromal cells and bone-marrow derived cells. However, proximal tubule cells appear to account for half or more of total renal mRNA. Despite limitations created by the presence of multiple cell types, bulk-kidney RNA-Seq can be interpretable; particularly when changes in cell-type specific markers are observed.

## METHODS

### Animals

2-month-old male C57BL/6 mice (Taconic, Hudson, NY) were maintained in standard conditions with free access to food and water. All animal experiments were conducted in accordance with NIH animal protocol H-0047R4.

### Microdissection

Mice were euthanized by cervical dislocation. The right kidney was rapidly removed and, after removal of the capsule, was immediately transferred to Trizol reagent for RNA extraction. The left kidney was placed in ice-cold dissection solution (135 mM NaCl, 1 mM Na_2_HPO_4_, 1.2 mM MgSO_4_, 5 mM KCl, 2 mM CaCl_2_, 5.5 mM glucose, 5 mM HEPES, 5mM Na acetate, 6mM alanine, 1mM trisodium citrate, 4mM glycine, 1mM heptanoate, pH 7.4) for microdissection. Cortical collecting ducts (CCDs), cortical thick ascending limbs (cTALs) and proximal tubule S2 segments (PTS2) were manually dissected in ice-cold dissection solution without protease treatment under a Wild M8 dissection stereomicroscope equipped with on-stage cooling. These segments are clearly identifiable because of its presence in the cortical medullary rays. After a thorough wash in ice-cold PBS (2 times), the microdissected tubules were transferred to Trizol reagent for RNA extraction. 1 to 4 tubules were collected for each sample.

### Whole-kidney RNA-Seq and single-tubule RNA-Seq

These steps were conducted as previously reported.^10^ Briefly, total RNA from whole kidney and microdissected proximal tubules were extracted using Direct-zol RNA MicroPrep kit (Zymo Research, Irvine, CA) and cDNA was generated by SMARTer V4 Ultra Low RNA kit (Clontech, Mountain View, CA) according to the manufacturer’s protocols. 1 ng cDNA was fragmented and barcoded using Nextera XT DNA Sample Preparation Kit (Illumina, San Diego, CA). Libraries were generated by PCR amplification, purified by AmPure XP magnetic beads, and quantified using a Qubit 2.0 Fluorometer. Library size distribution was determined using an Agilent 2100 bioanalyzer with a High-Sensitive DNA Kit (Agilent, Wilmington, DE). Libraries were pooled and sequenced (paired-end 50bp) on Illumina Hiseq 3000 platform to an average depth of 60 million reads per sample.

### Data processing and transcript abundance quantification

Data processing was performed as previously reported.^10^ Briefly, raw sequencing reads were processed by FASTQC (https://www.bioinformatics.babraham.ac.uk/projects/fastqc/) and aligned by STAR^33^ to the mouse Ensembl genome (Ensembl, GRCm38.p5) with Ensembl annotation (Mus_musculus.GRCm38.83.gtf). Unique genomic alignment was processed for alignment visualization on the UCSC Genome Browser. Transcript abundances were quantified using RSEM^5^ in the units of transcripts per million (TPM). Unless otherwise specified, the calculations were done on the NIH Biowulf High-Performance Computing platform.

### Whole kidney and proximal tubule transcriptomes

The mean TPM values were calculated across all samples: 3 mice, (whole kidney, n=3) and (S2 proximal tubule, n=12). These filtered data are reported on specialized publicly accessible, permanent web pages to provide a community resource: https://hpcwebapps.cit.nih.gov/ESBL/Database/MusRNA-Seq/index.html.

### Data deposition

The FASTQ sequences and metadata reported in this paper have been deposited in NCBI’s Gene Expression Omnibus (GEO) database, (accession number: GSE111837; https://www.ncbi.nlm.nih.gov/geo/query/acc.cgi?acc=GSE111837, secure token: crqzssqurzkbzsp).

### Curation of list of cell-type selective genes

To identify a list of cell-type selective genes from renal tubule segments, we used data from microdissected rat renal tubules published by Lee et al.^8^ as well as data from mouse microdissected tubules and single cells described by Chen et al.^10^ and Park et al.^15^ For other cell types, markers were determined using a combination of the following sources: general PubMed searches for publicly accessible research articles, commercial information sources for recommended marker antibodies, and general reference textbooks. Specific sources are given in Supplemental Dataset 3. The curated list was designed to be representative but not exhaustive.

## Acknowledgments

The work was primarily funded by the Division of Intramural Research, National Heart, Lung, and Blood Institute (project ZIA-HL001285 and ZIA-HL006129, M.A.K.). Next-generation sequencing was done in the National Heart, Lung and Blood Institute (NHLBI) DNA Sequencing Core Facility (Yuesheng Li, Director).

## Author Contributions

L.C., C.-L.C, H.J.J and M.A.K designed research; L.C., C.-L.C, and H.J.J performed research; J.Z.C. and M.A.K analyzed data; L.C., J.Z.C. and M.A.K wrote the paper.

---

**Supplemental Figure 1. Mapping quality of the whole-kidney RNA-Seq data.** Distribution of reads shows that uniquely mapped reads exceeds 85% of total reads in all three whole kidney samples. Total reads were: sample 1, 66142467; sample 2, 68482027; sample 3, 69079531.

**Supplemental Figure 2. Mapping quality of the microdissected proximal tubule S2 RNA-Seq data.** Distribution of reads shows that uniquely mapped reads exceeds 85% of total reads in all twelve S2 proximal tubule samples. Total reads were: sample 1, 69808466; sample 2, 84962667; sample 3, 75565121; sample 4, 74862689; sample 5, 76598350; sample 6, 78381995; sample 7, 70858077; sample 8, 77120838; sample 9, 64935558; sample 10, 69894298; sample 11, 70091668; sample 12, 67011247.

